# Neuronal *Ndst1* depletion accelerates prion protein clearance and slows neurodegeneration in prion infection

**DOI:** 10.1101/2023.06.19.545528

**Authors:** Patricia Aguilar-Calvo, Adela Malik, Daniel R. Sandoval, Christopher Barback, Christina D. Orrù, Heidi G. Standke, Olivia Thomas, Chrissa A. Dwyer, Donald P. Pizzo, Jaidev Bapat, Katrin Soldau, Ryotaro Ogawa, Mckenzie B. Riley, K. Peter R. Nilsson, Allison Kraus, Byron Caughey, Jeffrey J. Iliff, David Vera, Jeffrey D. Esko, Christina J. Sigurdson

## Abstract

Select prion diseases are characterized by widespread cerebral plaque-like deposits of amyloid fibrils enriched in heparan sulfate (HS), a major extracellular matrix component. HS facilitates fibril formation *in vitro*, yet how HS impacts fibrillar plaque growth within the brain is unclear. Here we found that prion-bound HS chains are highly sulfated, and that the sulfation is essential for HS accelerating prion conversion *in vitro*. Using conditional knockout mice to deplete the HS sulfation enzyme, Ndst1 (N-deacetylase, N-sulfotransferase), from neurons or astrocytes, we investigated how reducing HS sulfation impacts survival and prion aggregate distribution during a prion infection. Neuronal Ndst1-depleted mice survived longer and showed fewer and smaller parenchymal plaques, shorter fibrils, and increased vascular amyloid, consistent with enhanced aggregate transit toward perivascular drainage channels. The prolonged survival was strain-dependent, affecting mice infected with extracellular, plaque-forming, but not membrane bound, prion strains. Live PET imaging revealed rapid clearance of prion protein monomers into the CSF in mice expressing unsulfated HS, further suggesting that HS sulfate groups hinder transit of extracellular prion monomers. Our results directly show how a host cofactor slows the spread of prion protein through the extracellular space and identify an enzyme target to facilitate aggregate clearance.

**Author summary:** Prions cause a rapidly progressive neurologic disease and death with no curative treatment available. Prion aggregates accumulate exponentially in the brain in affected individuals triggering neuronal loss and neuroinflammation. Yet the additional molecules that facilitate aggregation are largely unknown, and their identification may lead to new therapeutic targets. We have found that prions in the brain preferentially bind to a highly sulfated endogenous polysaccharide, known as heparan sulfate (HS). Here we use genetically modified mice that express poorly sulfated neuron-derived HS, and infect mice with different prions strains. We find that the mice infected with a plaque-forming prion strain show a prolonged survival and fewer plaques compared to the controls. We also found that the prion protein was efficiently transported in the interstitial fluid in mice having poorly sulfated HS, suggesting that the prion protein is more readily cleared from the brain. Our study provides insight into how HS retains prion aggregates in the brain to accelerate disease and indicates the specific HS biosynthetic enzymes to target for enhancing protein clearance.

## Introduction

The spread of aberrant protein aggregates through the brain is a key pathogenic mechanism in Alzheimer’s, Parkinson’s, and prion disease^1–6^. Potential mechanisms for aggregate spread include bulk transport within the extracellular space (ECS)^7–11^. The ECS harbors a dynamic reservoir of interstitial fluid (ISF) that flows toward perivascular clearance pathways^11^. Evidence suggests that bulk ISF flow is enhanced during sleep, fostering the efflux of metabolic waste and proteins, such as amyloid-β (Aβ), from the brain^12–14^. Thus, alterations in the structure or composition of perivascular clearance pathways^15^ or the extracellular matrix (ECM)^16^ during aging or neurodegenerative disease could hinder the clearance of peptides and oligomers^17^.

Prion diseases are rapidly progressive neurodegenerative disorders in which prion protein aggregates (PrP^Sc^) deposit on cell membranes or as plaques embedded in the brain ECM, depending on the prion conformation^18–21^. Similar to Aβ plaques, PrP^Sc^ plaques are highly enriched in a prominent component of the ECM, HS^22–25^. Tissue HS proteoglycans (HSPGs) consist of long carbohydrate chains (40-300 alternating residues of glucuronic acid and glucosamine residues, also depicted as disaccharides) attached to one or more protein cores^26, 27^. A large body of evidence implicates HS in fibril formation and in the endocytosis of protein aggregates *in vitro*^28–32^. Moreover, polyanions administered intraventricularly increase survival time in experimental rodent models and in patients, potentially by blocking prion binding to endogenous HS^33–44^. Transgenic expression of mammalian heparanase, an enzyme that degrades HS, delays prion onset and progression^45^.

HS chains bind and concentrate proteins, such as growth factors and cytokines, and the level and pattern of sulfation determine the binding affinity^26, 27^. Here, we show that brain-derived prions selectively bind highly sulfated HS chains. To understand how the sulfation level modifies prion aggregate spread and disease progression, we use conditional *Ndst1* knock-out mice to reduce the sulfation of either neuron or astrocyte generated HS, and challenge mice with diverse prion strains. We found that mice with reduced neuronal HS sulfation survived longer when infected with the extracellular plaque-forming prion strain. The prion conformational properties were largely unaltered except for a reduction in PrP^Sc^ size, as fibrils were shorter and prion aggregates more soluble. Additionally, there were fewer parenchymal plaques, while vascular fibrils accumulated in the meninges. Finally, in positron emission tomography (PET) scan studies, radiolabeled soluble PrP^C^ underwent faster transport through the brain of mice expressing poorly sulfated neuronal HS, leading to increased PrP^C^ efflux into the spinal cord. Together, our studies identify how reducing HS sulfation extends survival, reduces parenchymal plaques, increases PrP^Sc^ solubility, and accelerates PrP clearance, demonstrating *NDST1* as a potential therapeutic target.

## Results

### Highly sulfated HS binds PrP^Sc^

To first establish the level and composition of HS molecules bound to PrP^Sc^ as compared to whole brain lysate, we performed liquid chromatography - mass spectrometry (LC/MS) analysis on whole brain lysate, comparing to the HS molecules bound to PrP^Sc^ from the same brain (the latter previously reported)^46^ (Fig 1A). For two prion strains (ME7 and mCWD), HS bound to PrP^Sc^ was 7-9% more sulfated (N-, 2-O-, and 6-O-sulfated) than HS in brain lysate (Fig 1B-D), indicating that more highly sulfated HS selectively binds PrP^Sc^. Prion-bound HS was particularly enriched in D2S0, which is a disulfated (N- and 2-O-sulfated) disaccharide (Fig 1C, S1 and S2 Tables). We repeated this experiment with cerebral cortex from human sporadic prion disease (sCJD)-affected brain. Consistent with PrP^Sc^ from mouse brain, the HS bound to PrP^Sc^ was more highly sulfated [prion-bound: 63% sulfation versus brain lysate: 53%] (Fig 1E-G). Additionally, PrP^Sc^ from sCJD-affected cortex and from familial prion disease-affected cerebellum (F198S mutation in *PRNP*) selectively bound the D0A6 disaccharide (Fig 1F, S1 Fig, S3 and S4 Tables).

**Fig 1.**
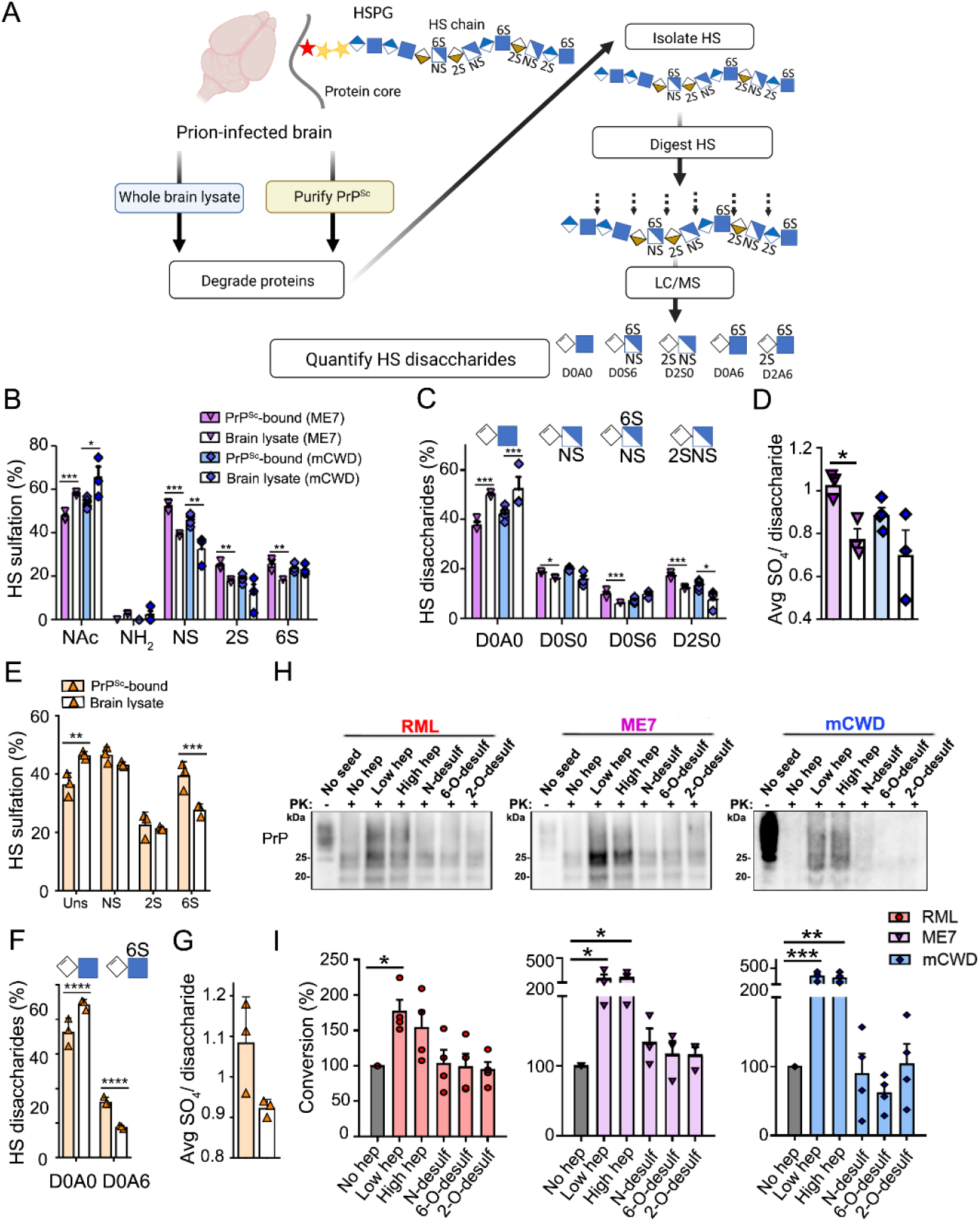
Mouse and human PrP^Sc^ bind highly sulfated HS. (A) Schematic of HS isolation from brain lysate or purified PrP^Sc^ for mass spectrometry analysis (a subset of the latter samples was previously published ^46, 79^). (B) Quantification of unsulfated (NAc and NH_2_) and sulfated (NS, 2S, and 6S) HS, (C) individual HS disaccharides, and (D) average sulfation per disaccharide of HS bound to PrP^Sc^ as compared to brain lysates (same brain) (*n* = 3 per strain). (E). Quantification of unsulfated and sulfated HS from sCJD-affected brain, (F) individual HS disaccharides, and (G) average sulfation per disaccharide of HS bound to PrP^Sc^ as compared to brain lysates (same brain). *N* = 3 per group (occipital cortex). (H) Representative western blots of PrP^Sc^-seeded PMCA in the presence or absence of heparin or heparin desulfated at positions N-, 2-O, or 6-O (No seed: no PrP^Sc^, No hep: no heparin, Low hep: 225 µg/ml of heparin, High hep: 2.225 mg/ml of heparin, N-, 6-O-, or 2-O-desulfated heparin: 225 µg/ml). (I) Quantification of PrP^Sc^ in PrP^Sc^-seeded PMCA experiments. N = 3 - 4 experimental replicates. Note for panels C and F, other disaccharides were not significantly different (shown in S1 - S3 Tables). \**P*< 0.05, \*\**P*< 0.01, \*\*\**P*< 0.005 and \*\*\*\**P*< 0.001, two-way ANOVA with Bonferroni’s post test comparing within a single strain (panels B, C, E, and F), unpaired two- tailed t-test with Bonferroni’s post test (panels D and G), and one-way ANOVA with Tukey’s post test (panel I). NAc: N-acetylglucosamine (GlcNAc); NH_2_: glucosamine (GlcNH_2_); NS: N- sulfated glucosamine (GlcNS); 2S: 2-O-sulfated glucuronic or iduronic acids (2-O-S); 6S: 6-O-sulfated glucosamine (6-O-S); PK : proteinase K.

Heparin is more sulfated than HS and promotes the fibril formation of PrP^Sc^, Aβ, tau, and α-synuclein *in vitro*^47–52^. To determine how the HS sulfate position impacts the interaction with PrP, we tested a library of selectively desulfated heparin molecules lacking the N-, 6-O-, or 2-O-sulfate group in a prion conversion assay known as protein misfolding cyclic amplification (PMCA)^53^. A PrP^C^-expressing cell lysate was seeded with mouse prions (three strains) with and without heparin. Heparin increased prion conversion by 60 - 350% (Fig 1H-I). Notably, eliminating any sulfate group, NS, 6S, or 2S, reduced conversion to baseline, indicating that sulfation *at all three positions* was necessary to accelerate prion conversion (Fig 1H-I).

HS binds ligands through electrostatic interactions between the anionic sulfate groups and cationic amino acids in ligands^26^. To identify the HS binding domain on PrP^C^, we performed alanine substitutions in three positively charged domains and an asparagine-rich domain (S2A Fig). Segments 23-27 and 101-110 were critical to PrP^C^ – HS binding, as lysine and arginine substitutions at either site markedly reduced binding (nearly 3-fold), while asparagine to alanine substitutions (segment 171-174) did not impact binding (S2B-D Fig). Together, these results suggest that the electrostatic interaction between the N-terminal lysine and arginine residues of PrP^C^ and the HS sulfate groups are essential for PrP - HS binding.

### Depleting neuronal HS sulfate groups reduces parenchymal prion plaques and prolongs survival

Given the findings that PrP^Sc^ selectively binds highly sulfated HS and that the sulfate groups enhance PrP conversion *in vitro*, we predicted that reducing HS sulfation would slow PrP conversion kinetics. To determine how less sulfated HS in the brain impacts PrP^Sc^ conformation and disease progression *in vivo*, we used *Ndst1^f/f^*mice to conditionally delete *Ndst1* (gene that encodes for the N-deacetylase/N-sulfotransferase enzyme that catalyzes the deacetylation and sulfation of HS chains at position N) from neurons by crossing mice to a neuron-specific Cre− recombinase line driven by the *synapsin1* promoter (*SynCre*) (S3A Fig). To validate the mouse line, an LC/MS analysis of the *Ndst1^f/f^SynCre^+/−^*and *SynCre^−/−^* (subsequently referred to as *SynCre+* and *SynCre−*) brain was performed and revealed that the *SynCre+* brain had less sulfated HS (33% reduced average sulfation per disaccharide) and less N-, 2-O- and 6-O-sulfated disaccharides, particularly D2S0 and D2S6 disaccharide units (S3B-C Fig, S5 Table). The *SynCre+* mice showed no clinical phenotype, brain lesions, nor change in PrP^C^ expression in the brain (S3D-H Fig), but showed lower microglial reactivity in cortex, thalamus, and hippocampus as compared to *SynCre−* mice (S3E-F Fig).

Sulfated HS enhances the uptake of prion fibrils in cultured cells^28, 54^ and may impact the conversion and spread of cell membrane-bound (GPI-anchored) and extracellular (GPI-anchorless) prions. Therefore, the *SynCre*+ and *SynCre^-^* mice were inoculated with prions that form either diffuse (RML; GPI-anchored) or diffuse and small plaque-like deposits (ME7; GPI-anchored and anchorless^46^). Mice infected with RML prions showed no difference in survival time [*SynCre−*: 158 ± 9 and *SynCre+*: 168 ± 10 days post-inoculation (dpi)] or brain lesions (S4A-C Fig), indicating that neuronal HS (nHS) sulfation level had no impact on the replication or spread of a GPI-anchored, oligomeric prion strain. In contrast, *SynCre*+ mice infected with ME7 prions showed a significantly prolonged survival [*SynCre−*: 172 ± 4 and *SynCre+*: 199 ± 11 dpi] (S4A Fig). Notably, the prolonged survival was not associated with differences in the spongiosis, glial activation, or PrP^Sc^ level or distribution in brain, nor in the PrP^Sc^ biochemical properties, including electrophoretic mobility and glycoprofile (S4B-H Fig), suggesting that reducing nHS sulfation primarily affected the kinetics of prion conversion and spread.

Since prion fibrils bind 4- to 10-fold more HS than subfibrillar aggregates (µg HS / µg PrP^Sc^)^46^, we predicted that reducing HS sulfation would substantially prolong survival in mice infected with the plaque-forming, GPI-anchorless prion strain, mCWD^55^. Because the disease course for mCWD prions in WT mice is more than 550 days, we crossed the *Ndst1^f/f^SynCre^+/−^* mice to *tga20^+/+^* mice, which overexpress PrP^C^ (4- to 6-fold) and show accelerated disease progression^55, 56^. Strikingly, mCWD-infected *Ndst1^f/f^tga20^+/+^SynCre^+/−^* mice (hereafter noted as *SynCre+*) displayed a markedly prolonged survival, approximately 40% longer than the *SynCre−* mice [*SynCre−*: 153 ± 19 and *SynCre+*: 206 ± 49 dpi] (Fig 2A-B).

**Fig 2.**
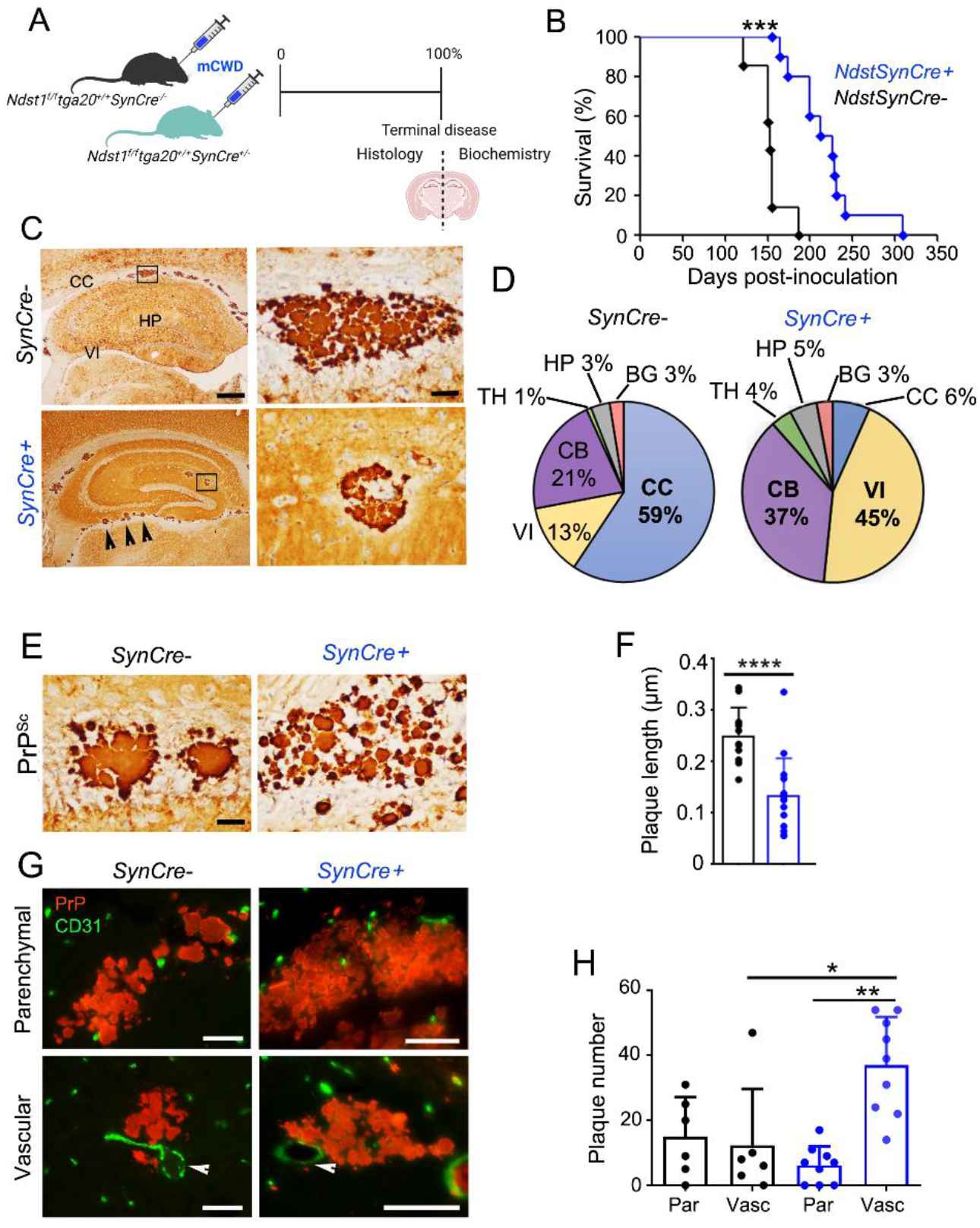
Reducing neuronal HS sulfation prolongs survival and lessens plaque load in prion-affected mice. (A) Schematic illustrates mCWD prion inoculation and tissue collection. (B) Survival curves for mCWD-infected *SynCre−* (*n* = 10) and *SynCre+* mice (*n* = 15). (C) Representative images reveal mCWD prion plaques in the corpus callosum (CC) of *SynCre−* mice and within vessels of the velum interpositum (VI) and hippocampus (HP) of *SynCre+* mice. Higher magnification depicted in right panels. Scale bars represent 500 μm (left) and 50 μm (right). (D) Pie charts show the plaque distribution in brain. VI and cerebellar (CB) aggregates were primarily vascular (meninges) (*n* = 6 *SynCre−* and 9 *SynCre+* mice). (E) Representative images of PrP^Sc^ immunolabelled plaques in CC. Scale bar = 50 μm. (F) Quantification of the plaque length in CC (from *n* = 33 and 38 plaques in *SynCre−* and *SynCre+*, respectively) (*n* = 4 mice per genotype). (G) Dual immunostaining of mCWD-infected brain sections for PrP^Sc^ and endothelial cells (CD31) (parenchymal plaques: corpus callosum; *SynCre−* vascular plaque: basal ganglia; *SynCre+* vascular plaque: thalamus). Scale bar represents 50 μm. (H) Quantification of parenchymal (par) and vascular (vasc) plaques throughout the brain. *N* = 6 *SynCre−* and 9 *SynCre+* mice. \**P*< 0.05, \*\**P*< 0.01, \*\*\**P*< 0.005, and \*\*\*\**P*< 0.001, Log-rank (Mantel-Cox) test (panel B), unpaired two-tailed t-test with Bonferroni’s post test (panel F), and two-way ANOVA with Bonferroni’s post test (panel H). TH thalamus, HT: hypothalamus, BG: basal ganglia, and CT: cerebral cortex.

Decreasing nHS sulfation had a pronounced effect on the prion plaque distribution in the *SynCre+* brain. While most *SynCre−* mice (83%; n = 5/6 mice) accumulated parenchymal plaques in the corpus callosum (CC), only one-third of the *SynCre+* developed plaques in the CC (33%; n = 3/9 mice) (Fig 2C-D), which were smaller and multicentric (clusters of small plaques) (Fig 2E-F). The *SynCre+* brain also showed an increase in vascular amyloid (*SynCre−:* 12 ± 2 versus *SynCre+:* 37 ± 2 amyloid-laden vessels), particularly within the velum interpositum and cerebellar meninges (Fig 2C-D, G-H and S5A-B Fig), suggestive of enhanced prion transit toward perivascular drainage pathways. There were no differences in the astrocytic or microglial response to aggregates (S5C-D Fig). Additionally, HS was still detectable in plaques (S5E Fig), and there was no change in plaque morphology in situ by electron microscopy (S5F Fig). Finally, fluorescent lifetime decay (FLIM) of a prion-bound fluorescent probe, heptameric formic thiophene acetic acid (h-FTAA), revealed no differences, suggesting no change in the PrP^Sc^ conformational properties (S6A Fig). Thus, reducing HS sulfation prolongs survival and reduces parenchymal plaques and plaque size in the brain, without detectably altering PrP^Sc^ secondary or tertiary conformation.

To further test PrP^Sc^ fibril conformation using biochemical analyses, we measured the biochemical properties of PrP^Sc^, including the electrophoretic mobility of the proteinase K (PK)-resistant core, glycoprofile, stability in chaotropes, and aggregate solubility (S6B-D Fig and Fig 3A-B). Notably the only difference was in PrP^Sc^ solubility, in which there was an increase (25%) in the *SynCre^+^*brains (Fig 3A-B) suggesting that the aggregate population was generally smaller. To further assess aggregate size, we next purified prion fibrils from brain homogenate and measured the length of isolated fibrils on electron microscopy grids. We found that fibrils were approximately 20% shorter (on average) in *SynCre+* brains (Fig 3C-D), consistent with a change in aggregate size as suggested by the higher PrP^Sc^ solubility.

**Fig 3.**
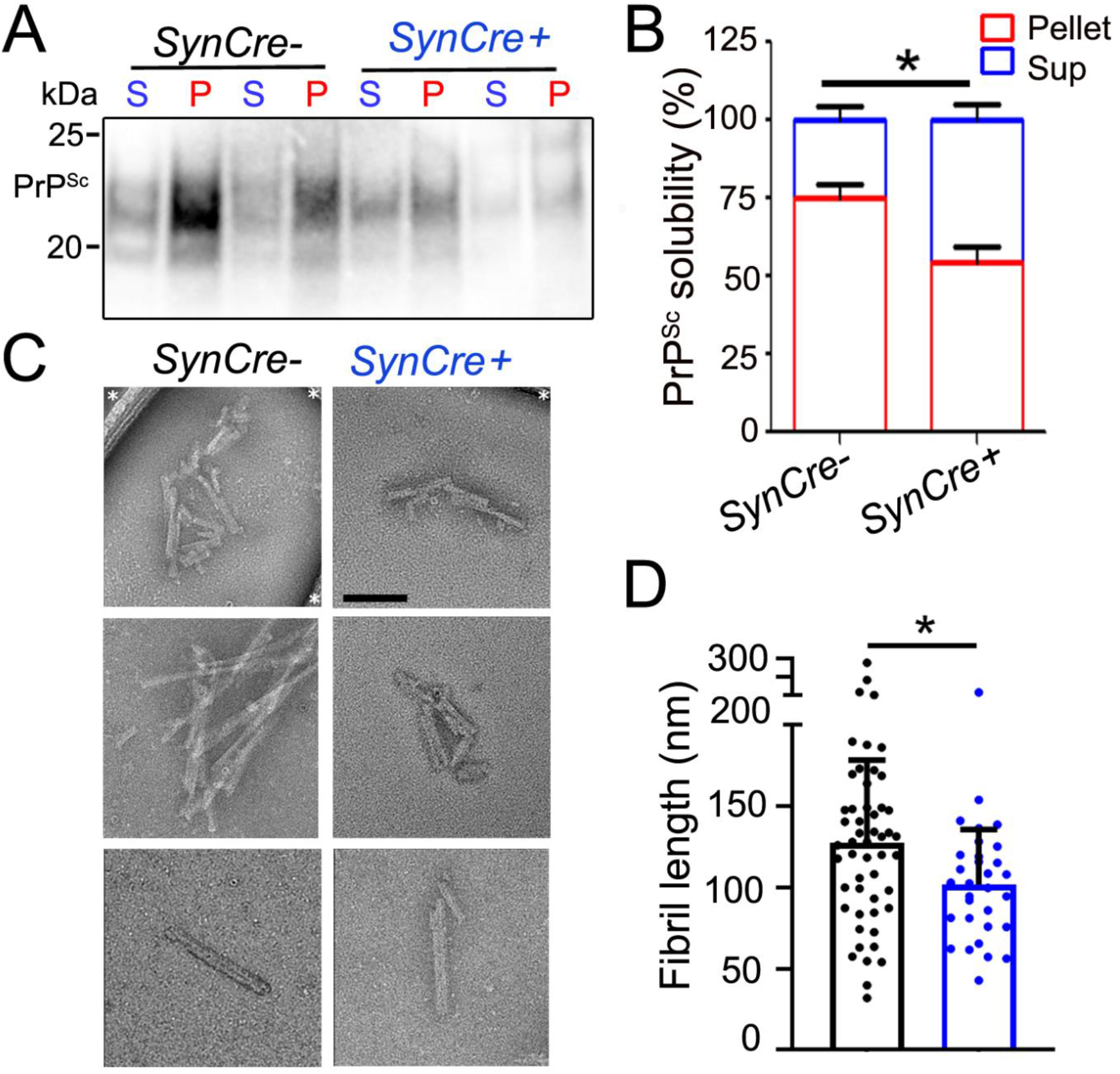
Reducing neuronal HS sulfation increases mCWD solubility. (A) Representative western blot of mCWD aggregates after PK digestion and centrifugation over an Optiprep layer (P = pellet, S = supernatant). (B) Quantification of PrP^Sc^ in pellet and supernatant fractions. *N* = 6 samples per genotype. (C) Representative electron microscopy images of purified PrP^Sc^ fibrils from mCWD-infected mice. Scale bar represents 100 nm. Asterisks indicate grid support film. (D) Quantification of the fibril lengths. *N* = 52 and 32 fibrils measured from *SynCre−* and *SynCre+* brains, respectively (*n* = 6 mice per genotype). \**P*< 0.05, unpaired two-tailed t-test with Bonferroni’s post test (panels B and D).

The increased PrP^Sc^ clearance into the cerebrospinal fluid (CSF) may lead to enhanced deposition in the spinal cord. Thus, we next compared the PrP^Sc^ seeding activity in the spinal cord from *SynCre*− and *SynCre*+ using real-time quaking-induced conversion (RT-QuIC) (S6 Table). However, there were no differences in the proportion of mice with seeding activity in spinal cord (*SynCre*−: 7 of 11 mice, *SynCre*+: 14 of 16 mice; Fisher’s exact test; *p* = 0.19), suggesting that reducing Nhs sulfation does not lead to increased incidence of PrP^Sc^ deposition in the spinal cord.

### Decreasing neuronal HS sulfation enhances PrP^C^ clearance into the spinal cord

Given that i) HS is a significant component of the brain ECM and ii) decreasing nHS sulfation reduces parenchymal plaque load and prolongs survival in prion-infected mice (Fig 2B-H and S5A Fig), we reasoned that HS may entrap PrP in the ECM, hindering clearance by ISF bulk flow. To investigate how reducing HS sulfation affects soluble PrP^C^ transit through the ISF, we used PET to track radiolabeled PrP in real time (Fig 4A). Recombinant PrP^C^ was radiolabelled with zirconium-89 (Zr89) (PrP-Zr89) (radiolabel confirmed in S7A Fig). Zr89-labelled PrP^C^ was stereotaxically injected into the left caudate putamen of *Ndst1^f/f^tga20^+/+^SynCre^+/−^* and *SynCre^−/−^* mice in two experiments. Mice were imaged by PET immediately after injection [day 0 (D0)] and at 20 hours post-injection (D1) (Fig 4B and S7 Fig).

**Fig 4.**
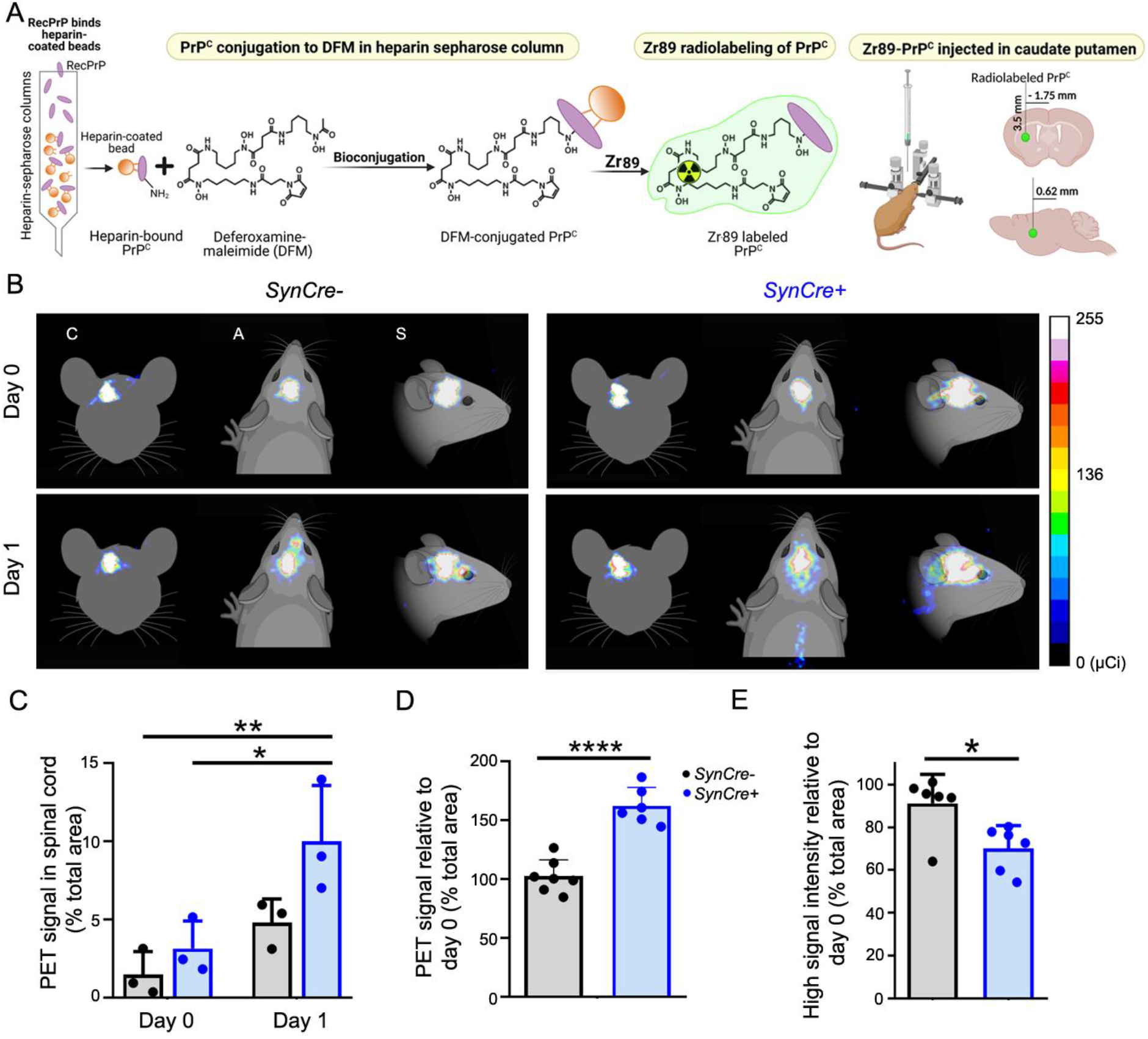
Rapid transit of Zr89-PrP^C^ through the brain and spinal cord of live *Ndst1^f/f^tga20^+/+^SynCre*+ mice. (A) Schematic shows the conjugation, radiolabeling, and stereotaxic injection of recPrP into anesthetized *Ndst1^f/f^tga20^+/+^SynCre*+ and *SynCre−* mice. (B) Representative PET scan images (coronal (C), axial (A) and sagittal (S) sections) of Zr89-PrP in *SynCre* and *SynCre+* mice immediately after PrP injection into the caudate putamen (day 0) and 20 hours later (day 1) in *n* = 6 animals per genotype. Graphs show (C) radioactive PrP in the spinal cord at day 0 and day 1 relative to the total PET signal area, as well as (D) the total PET signal area at day 1 relative to day 0 (sagittal) (experiments combined) and (E) the high PET intensity signal area (signal > 100 µCi) at day 1 relative to day 0 (panel C shows only Expt 1). \**P*< 0.05, \*\**P*< 0.01, and \*\*\*\**P*< 0.001, two-way ANOVA with Bonferroni’s post test (panel C), and unpaired two-tailed t-test with Bonferroni’s post test (panels D and E).sss

PET imaging immediately after injection revealed intense radioactive signal (more than 100 µCi) core similarly concentrated at the injection site in *SynCre*− and *SynCre*+ brains, with less than 5% in the spinal cord (Fig 4B-D and S7B-C Fig). Strikingly, at D1, the total PrP-Zr89 radiolabeled sagittal area had increased by 68% in the *SynCre*+ mice, in contrast to minimal change (9%) in the *Cre*− brains (Fig 4B, D and S7D Fig). Moreover, there was a pronounced decrease (30%) in the area occupied by the intense radioactive core (more than 100 μCi) in the *SynCre*+ mice (versus 9% in *Cre*−) (Fig 4E), suggestive of enhanced PrP-Zr89 diffusion from the injection site. Additionally, there was an approximately 14-fold increase in radioactivity in the spinal cord of *SynCre*+ as compared to *SynCre−* mice at D1 (*SynCre*−: 20% versus *SynCre*+: 287% increase compared to D0) (Fig 4C), indicating that more PrP-Zr89 had spread from the brain to the spinal cord in the *SynCre*+ mice as compared to *SynCre−* mice. Collectively these studies suggest that decreasing HS sulfation accelerates PrP^C^ clearance.

### Altering astrocytic HS sulfation does not impact prion disease progression

Since the HS in the CNS is secreted by neurons and glial cells^57–59^, we next tested how reducing astrocytic HS sulfation would affect prion conversion *in vivo*. *Ndst1^f/f^* mice were crossed with mice expressing *GFAPCre* (glial fibrillary acidic protein), as well as to *tga20* mice for mCWD inoculation. Brain lysates from uninfected aged *Ndst1^f/f^GFAPCre*+ (hereafter *GFAPCre*+) mice showed modest changes in HS composition (S8A-C Fig, S7 Table). Mice showed no change in PrP^C^ expression level, and no clinical or neuropathologic phenotype (S8D-F Fig). *GFAPCre*+ and *GFAPCre*− littermates were intracerebrally inoculated with RML, ME7, or mCWD prions. Interestingly, the prion-infected *GFAPCre*+ and *Cre*− showed similar survival times [Cre+ and Cre−: RML: 160 ± 16 versus 162 ± 5 dpi, ME7: 182 ± 14 versus 191 ± 10 dpi, mCWD: 140 ± 20 versus 133 ± 17 dpi, respectively]. Additionally, there were no differences in brain lesions (PrP^Sc^ distribution, vacuolation, or astrocytic gliosis) or biochemical properties of PrP^Sc^ (S8G-P Fig). These results indicate that reducing astrocytic HS sulfation had no detectable impact on prion disease progression, pathologic phenotype, or PrP^Sc^ biochemical properties for mice infected with three distinct strains of prions, in striking contrast to the effect of reducing neuronal HS sulfation.

In summary, our study demonstrates a previously unrecognized role for highly sulfated HS chains advancing prion disease progression. We found evidence that HS and PrP bind by electrostatic interaction, with PrP^Sc^ and sulfated HS engaging within the brain parenchyma, enhancing fibril elongation, parenchymal plaque formation, and markedly accelerating disease. Interestingly, this effect was cell source and prion strain dependent, as reducing neuronal, but not astrocytic HS, prolonged survival, and only impacted strains with a GPI-anchorless component. Finally, we systematically identified the more highly sulfated HS molecules concentrated within prion aggregates from mice and human brain by LC/MS, further supporting *NDST1* as a therapeutic target.

## Discussion

PrP^Sc^ and Aβ plaques are enriched in HS^23, 25, 46, 60^, yet whether HS facilitates fibril formation *in vivo* has been unclear. Here we establish that neuron-generated HS accelerates prion propagation in a strain-dependent manner. We show the importance and specificity of HS sulfate groups in binding PrP, as the HS molecules enriched within prion aggregates were more highly 6-O-, 2-O-, or N-sulfated compared to brain lysate, and sulfation was key to amplifying prion conversion. Notably, depleting nHS sulfation reduced parenchymal plaque load, expanded PrP^Sc^ deposition around meningeal vessels, and dramatically improved survival in mice infected with mCWD, collectively supporting a model of enhanced prion clearance in mice with poorly sulfated HS. Live imaging studies supported this model, revealing accelerated efflux of PrP monomer into the spinal cord. Thus, we propose that reducing sulfation of HS is sufficient to impair HS - PrP interaction and augment extracellular PrP egress into the CSF, prolonging the survival of mice with prion disease.

Previous studies report heparin and HS promote fibril formation *in vitro*^50, 61^, and prion-infected mice and vCJD patients treated with HS mimetics survive longer^33–35, 39–44^. Using genetic models, we and others recently found that shortening HS chains reduces plaque numbers as well as extends survival and improves behavior in prion-infected and Alzheimer’s mouse models, respectively^46, 62^, implicating HS in both prion and Aβ plaque formation *in vivo*. Our studies now demonstrate the highly sulfated nature of HS bound to prions using LC/MS, and the feasibility of manipulating HS biosynthetic pathways to enhance PrP clearance, increasing the resistance of mice to extracellular plaque formation. Notably, the few parenchymal plaques present in mice expressing less sulfated nHS were small and the isolated fibrils shorter and aggregates more soluble, together providing strong evidence to support that sulfated HS recruits prion subfibrillar aggregates, facilitating fibril elongation and plaque formation in the brain parenchyma. Whether other reported co-factors, such as lipids or nucleic acids^63–65^, also promote PrP conversion *in vivo* is unclear and would be important to address in future studies. Nevertheless, manipulating HS polymerase or sulfotransferase expression to reduce HS length or sulfation, respectively, may be a viable therapeutic strategy to promote the clearance of prions, Aβ, and other extracellular aggregates into the CSF.

Reducing nHS sulfation did not affect the disease progression of *Ndst1^f/f^SynCre^+/−^*mice infected with GPI-anchored prions (RML), despite PrP^C^ and PrP^Sc^ being embedded within an extensive meshwork of cell surface HSPGs. In vitro, HS reportedly promotes the endocytosis of PrP^Sc^, as well as tau and α-synuclein^28–31, 66, 67^, yet reducing nHS sulfation did not detectably alter RML prion propagation, astrocyte reactivity, or neurotoxicity. Although we found that heparin binds with high affinity and promotes GPI-anchored prion conversion *in vitro,* minimal HS was bound to GPI-anchored prions isolated from brain^46^, suggesting that the PrP^Sc^ membrane location limits access to HS, and HS may not enhance neurotoxicity in vivo. Given that GPI-anchored prions concentrate within lipid rafts, the membrane curvature or phosphate head groups, lipid raft size, or HS location and spatial orientation may hinder HS-PrP binding and constrain fibril elongation for membrane-bound prions, abrogating any major scaffolding effect by HS^68^. In contrast, extracellular GPI-anchorless PrP^C^ and PrP^Sc^ are unconstrained in the parenchyma and may more readily bind extracellular HS chains, consistent with our previous finding of abundant HS bound to extracellular prions^46^. That said, we cannot exclude that HS promotes the endocytosis, neuronal toxicity, or propagation of different prion conformers than used here, as prions propagating in different brain regions may access HS chains with a sulfation code better suited for binding.

Notably, only reducing nHS sulfation impacted disease progression; reducing astrocytic HS sulfation had no effect. Although both neurons and astrocytes produce and secrete HS^57, 58, 69^, PrP may selectively bind nHS due to the level and pattern of sulfation, or astrocytes may simply synthesize less HS. Alternatively, considering that only neuronal PrP^C^ is essential for PrP-linked toxicity^70^, it is conceivable that membrane-bound HSPGs such as syndecans or glypicans, facilitate extracellular PrP^Sc^ (GPI-anchorless) binding to PrP^C^ or neuronal surface receptor complexes in trans, potentiating neurotoxic signaling pathways. In this case, reducing nHS sulfation may decrease PrP^Sc^ interactions with cell surface receptors. Future studies may further elucidate the mechanism underlying the prolonged survival.

Our studies support a broader role of HS and the ECM in retaining proteins and promoting aggregation in the aging brain, with implications beyond prion disease. Given that HS also binds lipids and chemokines, this work suggests that increases in HS levels or sulfation may slow protein efflux and increase neuroinflammation, particularly in the presence of protein aggregates. Similar to PrP, the N-terminus of Aβ also binds to HS^71^ and HS sulfate groups are required for Aβ binding^72^. Given the similarities of our prion disease model to AD, future studies to (i) characterize how HS levels and composition change with age and disease in the human brain, (ii) determine whether HS enhances neuroinflammation, (iii) define the structure of HS concentrated within Aβ plaques, and (iv) devise strategies to enhance the clearance of Aβ in AD models by reducing HS sulfation would be of high priority.

We demonstrate a significant increase in the lifespan of prion-infected, nHS-depleted mice, which suggests that HS biosynthetic enzymes, such as Ndst1, may be therapeutic targets for select neurodegenerative diseases. Reducing HS sulfation in early disease would be expected to enhance aggregate transport toward perivascular clearance pathways. However, a caveat is a possible increase in meningeal vascular amyloid, as observed in prion-infected mice with reduced HS sulfation as shown here or with shortened HS chains^46^. Given that these results highlight the importance of HS sulfation in plaque formation, future work may define and inhibit PrP^Sc^ and Aβ binding to vessel-associated molecules to further promote prion clearance into the CSF.

In conclusion, our data provide evidence that PrP selectively binds highly sulfated neuronal HS, thereby identifying HS as the first integral *in vivo* co-factor in prion fibril formation and plaque assembly. Reducing HS sulfation decreased the parenchymal plaque burden and prolonged survival, suggesting that *in vivo*, PrP and HS electrostatic interactions are critical for binding. We also directly demonstrate how sulfated HS slows PrP^C^ clearance from the ISF in real time, indicating that GPI-anchorless PrP^C^ transits through the brain by bulk flow, and that clearance may be increased by manipulating HS chemical properties. Importantly, our data strongly supports the pursuit of therapeutic strategies targeting HS biosynthetic enzymes to facilitate protein aggregate clearance.

## Materials and Methods

### Prion transmission studies in mice

*Ndst1^f/f^* mice^73^ were bred to mice that express the Cre−recombinase under the neuron and astrocyte specific promoters, *synapsin1* and *glial fibrillary acidic protein* (*SynCre and GFAPCre*), respectively, and to *tga20* mice, which overexpress mouse PrP^C^^56^. Homozygosity for *tga20* was determined by quantitative real time PCR using the PureLink™ Genomic Purification Kit, and the Taqman™ Master Mix, Copy Number Assay (*Prnp*) and Copy Number Reference Assay, mouse, Tfrc (Thermo Fisher Scientific). Mice were maintained under specific pathogen-free conditions on a 12:12 light/dark cycle. All animal studies were approved by the Institutional Animal Care and Use Committee at UC San Diego. Protocols were performed in strict accordance with good animal practices, as described in the Guide for the Use and Care of Laboratory Animals published by the National Institutes of Health.

Male and female *Ndst1^f/f^SynCre+ or GFAPCre+* and *Ndst1^f/f^tga20^+/+^SynCre+ or GFAPCre*+ (6-8 weeks old), and *Cre−* littermate control mice, were anesthetized with ketamine and xylazine and inoculated into the left parietal cortex with 30 µl of 1% prion-infected brain homogenate prepared from terminally ill mice (n= 7 – 17 mice/group) or with 1% mock brain homogenate (n= 3 – 4 mice/group). Prion-inoculated mice were monitored three times weekly for the development of terminal prion disease, including ataxia, hyperactivity, kyphosis, stiff tail, hind leg clasp, and hind leg paresis, and were euthanized at the onset of terminal disease. During necropsy, the brain was halved, and the left hemisphere was immediately fixed in formalin. Fixed brains were treated for 1 hour in 96% formic acid, post-fixed in formalin, cut into 2 mm transverse sections, and paraffin-embedded for histological analysis. A 2-3 mm transverse section was removed from the left hemisphere at the level of the hippocampus/thalamus, embedded in optimal cutting temperature (OCT) compound and immediately frozen on dry ice. The remaining brain tissue was frozen for biochemical studies. Survival time was calculated from the day of inoculation to the day of terminal clinical disease.

### Histopathology and immunohistochemical stains

Five-micron sections were cut onto positively charged silanized glass slides and stained with hematoxylin and eosin (HE), or immunostained using antibodies for PrP (SAF84, epitope in the globular domain at the amino acids 160–170 of the mouse PrP), astrocytes (glial fibrillary acidic protein, GFAP), and microglia (Iba1). GFAP (DAKO; 1:6000), Iba1 (Wako; 1:3000), and PrP (Cayman Chemical; 1:1200) immunohistochemistry were performed on an automated tissue immunostainer (Ventana Discovery Ultra, Ventana Medical Systems, Inc). Each epitope had independently optimized retrieval parameters to yield the maximal signal to noise ratio. For PrP, slides were incubated in protease 2 for 20 minutes followed by antigen retrieval in CC1 (tris-based; pH 8.5; Ventana) for 64 minutes at 95 °C. For GFAP only the protease P2 was used (Ventana) for 16 minutes. Iba1 retrieval consisted of CC1 for 40 minutes at 95 °C. Following retrieval, antibodies were incubated on the tissue for 32 minutes at 37 °C. The secondary antibody (HRP-coupled goat anti-rabbit or anti-mouse; OmniMap system; Ventana) was incubated on the sections for 12 minutes at 37 °C. The primary antibody was visualized using DAB as a chromogen followed by hematoxylin as a counterstain. Slides were rinsed, dehydrated through alcohol and xylene and cover slipped. For the PrP and CD31 (endothelial cells) dual immunolabelling, tissue sections were stained sequentially using anti-PrP SAF84 (1:150) and CD31 antibodies (Dianova; 1:150) using the tyramide signal amplification system (TSA; ThermoFisher). Slides were stained on a Ventana Discovery Ultra (Ventana Medical Systems, Tucson, AZ, USA). Antigen retrieval was performed using a slightly basic treatment solution (CC1; pH 8.5, Ventana) for 92 minutes at 95 °C. Sections then were incubated in anti-PrP antibody for 32 minutes at 37 °C, followed by anti-mouse-HRP (UltraMap Detection Kit, Ventana) and TSA-Alexa 594. The antibodies were denatured by treatment in a citric acid-based solution, pH 6 (CC2, Ventana) for 24 minutes at 95 °C. Subsequently, the slides were incubated with anti-CD31 antibody (rat) for 32 minutes at 37 °C followed by rabbit anti-rat (1:500; Jackson ImmunoResearch) and detected using the OmniMap system (Ventana) to fluorescently label CD31-expressing cells with TSA-Alexa 488.

The PrP and HS dual immunolabelling was performed as previously described but with minor changes^46^. Briefly, tissue sections were deparaffinized, and epitope exposure was performed using formic acid, PK, and heated citrate buffer (pH 6). Sections were blocked and incubated with anti-PrP SAF-84 (1:400) antibody for 1 hour followed by IgG –CY3 (Jackson Immunolabs; 1:200) for 30 minutes. Sections were next incubated with anti-HS 10E4 antibody (AMS Bioscience; 1:300), anti-mouse IgM biotin (Jackson Immunolabs; 1:500) for 30 minutes, streptavidin-HRP (Jackson Immunoresearch; 1:2000) for 45 minutes and tyramide-Alexa488 (Invitrogen) for 10 minutes. Nuclei were labeled with DAPI, and slides were mounted with fluorescent mounting medium (ProLong^TM^ Gold antifade reagent). As controls for the HS stain, a subset of duplicate slides was treated with 8 milliunits of heparin lyases I, II and III for one hour prior immunostaining. Isotype immunoglobulin controls, single sections immunostained for PrP or HS, and prion negative (uninfected) cases were also included.

### Lesion profile

Brain lesions from prion-infected mice were scored for the level of PrP immunological reactivity, spongiosis, and gliosis on a scale of 0–3 (0= not detectable, 1= mild, 2= moderate, 3= severe) in 9 regions: (1) dorsal medulla, (2) cerebellum, (3) hypothalamus, (4) medial thalamus, (5) hippocampus, (6) septum, (7) medial cerebral cortex dorsal to hippocampus, (8) cerebral peduncle, and (9) cerebellar peduncle. A sum of the three scores resulted in the value obtained for the lesion profile for the individual animal in a specific brain area and was depicted in the ‘radar plots’. Two investigators blinded to animal identification performed the histological analyses.

### Quantitative analysis of astrocytic and microglial inflammation

To measure astrocytic gliosis and microglial activation in *Ndst1^f/f^SynCre, Ndst1^f/f^tga20^+/+^SynCre*, *Ndst1^f/f^GFAPCre* and *Ndst1^f/f^tga20^+/+^GFAPCre* mice, slides containing cerebral cortex, corpus callosum, hippocampus, thalamus, hypothalamus, and cerebellum were imaged using the Olympus EX41 microscope with DP Controller. Images were converted to grayscale, and FIJI (an ImageJ based image processing software) was used to measure the total brain area and quantify astrocyte and microglia reactivity using the “Measure” function. Astrocyte and microglia were demarcated using the “Find the edges” function and particle analysis was used to measure the area occupied. The total area covered by astrocyte and microglia was divided by the total area for each brain region.

### Western blot and glycoprofile analyses

For ME7 strain in WT mice, PrP^Sc^ was concentrated from 10% brain homogenate in phosphate buffered saline (PBS) (w/v) by performing sodium phosphotungstic acid (NaPTA) precipitation prior to western blotting^74^. Briefly, 20 μl of 10% brain homogenate in an equal volume of 4% sarkosyl in PBS was nuclease digested (benzonase^TM^, Sigma) followed by digestion with 20 μg/ml PK at 37 °C for 30 minutes. After addition of 4% sodium phosphotungstic acid in 170 mM MgCl_2_ and protease inhibitors (Complete-TM, Roche), extracts were incubated at 37 °C for 30 minutes and centrifuged at 18,000 x g for 30 minutes at 25 °C. Pellets were resuspended in 2% sarkosyl prior to electrophoresis and immunoblotting. Samples were electrophoresed in 10% Bis-Tris gel (Invitrogen) and transferred to nitrocellulose by wet blotting. Membranes were incubated with monoclonal antibody POM19 [discontinuous epitope at C-terminal domain, amino acids 201–225 of the mouse PrP^75^] followed by incubation with an HRP-conjugated IgG secondary antibody. The blots were developed using a chemiluminescent substrate (Supersignal West Dura ECL, ThermoFisher Scientific) and visualized on a Fuji LAS 4000 imager. Quantification of PrP^Sc^ glycoforms was performed using Multigauge V3 software (Fujifilm). For mCWD, 100 μl of 10% brain homogenate was concentrated using NaPTA as described above and digested with 100 μg/ml PK at 37 °C for 45 minutes.

### Conformation stability assay

To measure prion strain stability in guanidine chloride (GdnHCl), 10% brain homogenates were denatured for 1 hour in increasing concentrations of GdnHCl from 0 to 6 M. Samples were then diluted with a Tris-based lysis buffer (10 mM Tris-HCl, 150 mM NaCl, 10 mM EDTA, 2% sarkosyl, pH 7.5) to 0.15 M GdnHCl and digested with PK at a ratio of 1:500 (1 μg PK: 500 μg total protein) for 1 hour at 37 °C. The digestion was stopped with 2 mM phenylmethylsulfonyl fluoride (PMSF) and protease inhibitors (Complete-TM, Roche) followed by centrifugation at 18,000 x g for 1 hour. Pellets were washed in 0.1 M NaHCO_3_ (pH 9.8) and centrifuged at 18,000 x g for 20 minutes. Pellets were then denatured in 6 M guanidine isothiocyanate, diluted with 0.1 M NaHCO_3_, and coated passively onto an ELISA plate. PrP was detected with biotinylated-POM1 antibody (epitope in the globular domain, amino acids 121–231 of the mouse PrP^75^), a streptavidin HRP-conjugated secondary antibody, and a chemiluminescent substrate. Stability was measured in a minimum of 3 independent experiments, comparing *Ndst1^f/f^tga20^+/+^SynCre+* with *SynCre*– mice (3 – 4 mice per group).

### PrP^Sc^ solubility assay

Brain homogenates were solubilized in 10% sarcosyl in PBS and digested with 50 μg/ml of PK (final concentration) at 37 °C for 30 minutes. Protease inhibitors were added (Complete-TM, Roche), and samples were layered over 15% Optiprep™ and centrifuged at 18,000 x g for 30 minutes at 4 °C. Supernatants were removed and pellets were resuspended in PBS in a volume equivalent to the supernatant. Supernatant and pellet fractions were immunoblotted using anti-PrP antibody POM19. PrP signals were captured and quantified using the Fuji LAS 4000 imager and Multigauge V3.0 software. Brain samples were measured from 3-4 mice per genotype.

### Cell-lysate protein misfolding cyclic amplification (clPMCA)

The pcDNA3.1 vector (Invitrogen) with the mouse *Prnp* encoding the 3F4 epitope (109M, 112M human numbering) was used as a template for site-directed mutagenesis (QuikChange Site Directed Mutagenesis kit™) (Agilent). PrP-deficient RK13 cells (ATCC) were transfected with 5-10 µg of plasmid DNA using lipofectamine 3000 (Invitrogen). At 24 hours post-transfection, cells were washed twice in PBS, harvested in 1 ml PBS, and centrifuged for 1 minute at 1,000 x g. The pellet was resuspended in PMCA buffer (PBS containing 1% triton X-100, 150 mM NaCl, and 5 mM EDTA plus Complete-TM protease inhibitors), passed repeatedly through a 27-gauge needle, and clarified by centrifuging at 2000 x g for 1 minute.

RML, ME7, and mCWD prions were used to seed mouse PrP^C^ (no heparin reactions). The PrP^C^ was newly prepared for each independent experiment. The prion seeds were derived from brain homogenate that was pooled from mice inoculated with the same prion strain. The brain homogenate samples pooled to generate the seeds were consistent between the experiments. Prion-infected brain homogenate (10% w/v) was added to PrP^C^-expressing RK13 cell lysate (1:10, PrP^Sc^ : PrP^C^ by volume) and subjected to repeated 5 seconds sonication pulses (S4000, QSonica) with 10 minutes of incubation between each pulse, over a total period of 24 hours. Sonication power was maintained at 50-60% and samples were continuously rotated in a water bath at 37 °C. Samples were then digested with 200 µg/ml PK for 30 minutes at 37 °C and analyzed by western blot using the anti-PrP monoclonal antibody 3F4^76^. PrP^C^ levels were measured by blotting 1-2 µl from unseeded lysates. Signals were quantified using a Fujifilm LAS-4000 imager and Multi Gauge software and compared by percent conversion to control samples (considered 100%). PK-digested unseeded lysates were included in all experiments to exclude PrP^Sc^ contamination of the PMCA substrates and spontaneous assembly of mutant PrP^C^ protein. At least three independent experimental replicates were performed for each mutant and each prion strain used as seed. For the PCR assays with heparin (Scientific Protein Laboratories) and desulfated heparin (Tega), 1 µl of 225 µg/ml or 2,225 µg/ml sulfated heparin, or 1 µl of 225 µg/ml desulfated heparin (N-,6-O-, or 2-O-desulfated) were added to the PMCA reactions. The level of PrP^Sc^ formed in the presence and absence of heparin was measured by western blot.

### h-FTAA staining and fluorescence life time imaging

Sections (10 µm) of OCT-embedded brain samples were cut onto positively charged silanized glass slides, dried for 1 hour and fixed in 100% then 70% ethanol for 10 minutes each. After washing with deionized water, sections were equilibrated in PBS, pH 7.4, for 10 minutes. Heptamer-formyl thiophene acetic acid (h-FTAA; 1.5 mM in de-ionized water) was diluted in PBS to a final concentration of 1.5 μM and added to the sections. The sections were incubated with h-FTAA for 30 minutes at room temperature, washed with PBS, and mounted using Dako fluorescence mounting medium. The fluorescence decay of h-FTAA bound to PrP aggregates was collected using an inverted Zeiss (Axio Observer.Z1) LSM 780 microscope (Carl Zeiss MicroImaging GmbH) equipped with a modular FLIM system from Becker and Hickl. In this setup, the emitted photons were routed through the direct coupling confocal port of the Zeiss LSM 780 scanning unit and detected by a Becker and Hickl HPM-100-40 hybrid detector. Data was recorded by a Becker and Hickl Simple-Tau 152 system (SPC-150 TCSPC FLIM module) with the instrument recording software SPCM version 9.42 in the FIFO image mode, 256 × 256 pixels, using 256 time channels (Becker and Hickl GmbH). For all acquisitions, a T80R20 main beam splitter was used, and the pinhole was set to 20.2 μm. A 490 nm laser line from a pulsed tunable In Tune laser (Carl Zeiss MicroImaging GmbH) with a repetition rate of 40 MHz was used for excitation. Data was subsequently analyzed in SPCImage version 3.9.4 (Becker and Hickl GmbH), fitting each of the acquired decay curves to a tri-exponential function and color-coded images, as well as distribution histograms, showing the intensity-weighted mean lifetimes generated with the same software. The procedure of staining and FLIM imaging protein aggregates with h-FTAA is described in detail in reference {Nyström, 2017 #15836}.

### Purification of PrP^Sc^ for mass spectrometry and electron microscopy

To analyze HS bound to PrP^Sc^, PrP^Sc^ was first purified from mouse brains as previously described^46, 77^. Briefly, one ml of 10% brain homogenate was mixed with an equal volume of TEN(D) buffer (5% sarkosyl in 50 mM Tris-HCl, 5 mM EDTA, 665 mM NaCl, 0.2 mM dithiothreitol, pH 8.0), containing complete TM protease inhibitors (Roche). Samples were incubated on ice for 1 hour and centrifuged at 18,000 x g for 30 minutes at 4 °C. All but 100 μl of supernatant was removed, and the pellet was resuspended in 100 μl of residual supernatant and diluted to 1 ml with 10% sarkosyl TEN(D). Each supernatant and pellet was incubated for 30 minutes on ice and then centrifuged at 18,000 x g for 30 minutes at 4 °C. Supernatants were recovered while pellets were held on ice. Supernatants were added separately into ultracentrifuge tubes with 10% sarcosyl TEN(D) buffer containing protease inhibitors and centrifuged at 150,000 x g for 2.5 hours at 4 °C. Supernatants were discarded while pellets were rinsed with 100 μl of 0.25 M NaCl in TEN(D) buffer with 1% sulfobetaine (SB 3–14) and protease inhibitors and then combined and centrifuged at 200,000 x g for 2 hours at 20 °C. The supernatant was discarded, and pellet was washed and then resuspended in 200 μl of ice cold TMS buffer (10 mM Tris-HCl, 5 mM MgCl_2_, 100 mM NaCl, pH 7.0) with protease inhibitors. Samples were incubated on ice overnight at 4 °C. Using syringe and blunt needles, samples were homogenized and then incubated with 25 units/ml nuclease (benzonase^TM^, Sigma-Aldrich) and 50 mM MgCl_2_ for 30 minutes at 37 °C at 120 x g followed by a digestion with 1 mg/ml PK (final concentration) for 1 hour at 37 °C at 120 x g. PK digestion was stopped by incubating samples with 2 mM PMSF on ice for 15 minutes. Samples were incubated with 2 mM EDTA for 15 minutes at 37 °C at 120 x g. NaCl (0.25 M final) was added to all tubes followed by an equal volume of 2% SB 3–14 buffer. For the sucrose gradient, a layer of 0.5 M sucrose, 100 mM NaCl, 10 mM Tris-HCl, and 0.5% SB 3–14, pH 7.4 was added to ultracentrifuge tubes. Samples were then carefully transferred, and the tubes topped with TMS buffer. Samples were centrifuged at 200,000 x g for 2 hours at 20 °C. The pellet was rinsed with 0.5% SB 3–14 in PBS. Pellets were resuspended in 50 μl of 0.5% SB 3–14 in PBS and stored at −80 °C. Gel electrophoresis and silver staining were performed to assess the purity of PrP^Sc^.

### Heparan sulfate purification and analysis by mass spectrometry

Heparan sulfate (HS) was extracted from whole brain homogenates and purified by anion exchange chromatography, as previously described^46^. For depolymerization, HS chains were extensively digested with 1 milliunit each of heparin lyases I, II, and III (AMS Biotechnology). The disaccharides resulting from enzymatic depolymerization were tagged by reductive amination with [^12^C_6_] aniline and mixed with [^13^C_6_] aniline-tagged disaccharide standards. Samples were analyzed by liquid chromatography-mass spectrometry (LC-MS) using an LTQ Orbitrap Discovery electrospray ionization mass spectrometer (ThermoFisher Scientific). The disaccharides measured were: D0H0, D0A0, D0H6, D2H0, D0S0, D0A6, D2A0, D2H6, D0S6, D2S0, D2A6 and D2S6.

### Negative stain electron microscopy

400 mesh lacey carbon grids (Ted Pella) were glow discharged, placed on a 7 μl droplet of purified PrP^Sc^ sample, and incubated for 20 minutes in a humidified chamber. Grids were then blotted on filter paper, immersed briefly in Nano-W stain (Nanoprobes) and blotted again before incubating on a droplet of Nano-W for 1 minute. Grids were then blotted dry and imaged on a FEI Tecnai TF20 (200kV, FEG) with a 4k x 4k CMOS-based Tietz TemCam-F416 camera. Fibrils purified from terminally ill mCWD-infected *Ndst1^f/f^tga20^+/+^SynCre−* and *SynCre+* brains were distributed on the electron microscopy grids as single filaments as well as clusters of fibrils. Fibril lengths were assessed by three blinded investigators, and lengths recorded using Image J or iMOD software packages when both ends of a non-overlapping filament were readily visualized. [

### PrP^C^ mutant generation and heparin sepharose chromatography

To identify the HS-binding domains, three clusters of lysine- and arginine-residues and one cluster of asparagine residues were exchanged for alanine within mouse *Prnp* in a pcDNA3.1 vector by site-directed mutagenesis (QuikChange site-directed mutagenesis kit; Agilent). PrP-deficient RK13 cells (ATCC) were transfected with 10 μg of plasmid DNA using Lipofectamine 3000 (Invitrogen). At 24 hours post-transfection, cells were washed twice in PBS and digested with 0.75 milliunits of phospholipase C from *Bacillus cereus* (Sigma Aldrich) in 1.5 ml of Opti-MEM media in PBS (1:2 dilution) (ThermoFisherScientific) for one hour at 37 °C. The media was recovered and clarified by centrifugation at 2000 x g for 1 minute. Supernatants from the duplicate plates were pooled and saved for chromatography analysis.

For affinity chromatography, Heparin Sepharose 6 Fast Flow beads (Healthcare Life Sciences) were loaded into Bio-Spin® chromatography columns (Bio-Rad) and packed with 2 ml of equilibration buffer (0.15 M NaCl in 25 mM HEPES, pH 7.4). Supernatants containing GPI-cleaved proteins, including WT and mutant PrP^C^, were applied onto the columns. The flow through was recovered, recirculated onto the column two times, and saved. The column was next washed with 2 ml of 0.15 M NaCl in 25 mM HEPES buffer (pH 7.4) and the unbound proteins were recovered in a clean tube. The bound PrP^C^ was step-eluted with 1 ml of elution buffer containing increasing concentrations of NaCl (300 mM – 2 M) in 25 mM HEPES. The unbound PrP^C^ in the 0.15 M wash and PrP^C^ in all eluates were analyzed for PrP^C^ level by immunoblot using POM19 antibody. At least three experimental replicates were performed for each PrP^C^ construct.

### RT-QuIC analysis of mCWD-inoculated mice

The RT-QuIC reaction mix was composed of 10 mM phosphate buffer (pH 7.4), 130 mM NaCl, 0.1 mg/ml recombinant Syrian golden hamster prion protein (residues 90-231; rPrP^Sen^), 10 μM thioflavin T (ThT), 1 mM ethylenediaminetetraacetic acid tetrasodium salt (EDTA), and 0.002% SDS. Each well of a black 96-well plate with a clear bottom (Nunc) was loaded with aliquots of reaction mix (98 μl) and seeded with 2 μl of a 10^-2^ to 10^-4^ dilution of 10% mCWD spinal cord homogenate. The plate was sealed (plate sealer film, Nalgene Nunc International), incubated at 50°C in a BMG FLUOstar Omega plate reader and subjected to cycles of 1 min shaking (700 rpm double orbital) and 1 min rest with ThT fluorescence measurements (450 +/− 10 nm excitation and 480 +/− 10 nm emission; bottom read) taken every 45 minutes. Reactions were classified as RT-QuIC positive based on a threshold set at 10% of the maximum ThT fluorescence value on each plate.

### PrP^C^ conjugation to deferoxamine-maleimide

To label PrP with zirconium-89, full length recombinant mouse PrP (23-230) generated in *E.coli* was first conjugated to deferoxamine-maleimide (Macrocyclics) to produce deferoxamine-conjugated PrP^C^ (DFO-PrP^C^). To ensure that linker was not conjugated to the lysine-rich heparin binding domain, PrP was bound to heparin sepharose beads to block the heparin binding sites. To do this, Heparin Sepharose 6 Fast Flow beads (1 ml) (Healthcare Life Sciences) were loaded into disposable Bio-Spin® chromatography columns (Bio-Rad) and packed with 2 ml of equilibration buffer (0.15 M NaCl in 25 mM HEPES, pH 7.4). Recombinant PrP (200 µg) was mixed with 1 ml of wash buffer [0.15 M NaCl, 25 mM HEPES] and applied onto the columns. The columns were washed with 3 ml of wash buffer. The beads were then transferred to a 1.5 ml eppendorf polypropylene tube and incubated with 200 µl of DFM for 24 hours at room temperature with rotation. The bead slurry was transferred to a chromatography column and the PrP conjugation was stopped after 24 hours by adding 600 µl PBS with 0.1 M glycine (pH 7). The beads were washed with 2 ml equilibration buffer, and the conjugated PrP (PrP-DFM) was eluted with 1 ml of elution buffer (0.7 M NaCl, 25 mM HEPES, pH 7.2). PrP-DFM was concentrated using Zeba spin desalting columns.

### DFO-PrP radiolabeling with zirconium-89

Zirconium-89 (Zr89) oxalate (Washington University) adjusted to pH 7.5 with 0.5 M HEPES/ Na_2_CO_3_ (2 M) solution was incubated with 30 μg deferoxamine-conjugated PrP at room temperature for 60 minutes and diluted to approximately 100 μCi per injection following previously published procedures^78^. To measure the free Zr89 that has not been chelated, quality control was performed by instant thin layer chromatography (silica gel; 0.1 M citrate buffer pH 4.5); typical radiochemical purities (RF = 0.8) of 95 – 99 % were achieved. Typical specific volume was 25 μCi/μl. To confirm that PrP remained radiolabeled, Zr89-PrP^C^ at different dilutions was loaded in 10% Bis-Tris gel (Invitrogen) and electrophoresed four days after radiolabeling. Zr89 was added to the protein ladder (Precision Plus Protein Standard, Dual Color, Bio-Rad) at the expected size for recPrP, 25 kDa, and the gel was scanned in a phosphoimager (Typhoon).

### Stereotaxic injection of conjugated PrP

Mice (14-16 weeks old) (n = 3 - 4 mice/genotype/experiment) were anesthetized with isoflurane. Mice were weighed and placed in a three-point stereotaxic apparatus (Stoelting). A 2 cm midline incision was made in the skin over the sagittal suture to expose bregma, and a burr hole was drilled in the left parietal bone (0.62 mm caudal, −1.75 mm lateral to bregma) using an Ideal Micro-drill. A 22-gauge needle (Hamilton) was inserted to a depth of 3.5 mm and 1 µl of Zr89-PrP^C^ was injected at a rate of 75 nl/minute over 15 minutes using a Quintessential Stereotaxic Injector (Stoelting). The needle remained in the injection site for 10 minutes post-injection prior to removal from the brain to prevent backflow. Animals were next removed from the stereotaxic device and placed on the PET scanner (G.E. Vista), and radioactivity measurements were collected as dynamic scans using list mode over 30 minutes. Scans were repeated 20 hours later. PET images were reconstructed using Vista DR (G.E. Healthcare) software and the area and volume covered by radioactivity as well as the signal intensity were assessed with FIJI (an ImageJ based image processing software).

### Statistics

Log-rank (Mantel-Cox) tests were performed to assess survival differences between groups. A Student’s t-test (two-tailed, unpaired) with Bonferroni’s post test was used to determine the statistical significance between the *Ndst1^f^*^/*f*^*SynCre^+/−^*and *SynCre^−/−^* and *Ndst1^f^*^/*f*^*GFAPCre^+/−^*and *GFAPCre^−/−^* mouse groups for the PrP^C^ level of expression, lesion profiles, activated microglia, PrP^Sc^ glycoprofiles, PrP^Sc^ conformation stability and PrP^Sc^ fibril structure. One-way ANOVA with Tukey’s post test was performed to determine statistical significance in the levels of prion conversion by PMCA. Two-way ANOVA with Bonferroni’s post test was used to compare the composition of HS associated with different prion strains, the number of vascular versus parenchymal plaques, the binding affinity of PrP^C^ mutants with heparin, the levels of PrP in the spinal cord, and the area covered by high PET intensity signal (signal > 100 µCi). Unpaired two-tailed t-test with Bonferroni’s post test was used to compare the average sulfate groups per disaccharide as well as the plaque length and the mCWD fibril length and solubility in *Ndst1^f^*^/*f*^*SynCre^+/−^*and *SynCre^−/−^* brain. The proportion of *Ndst1^f/f^SynCre^+/−^* versus *SynCre^−/−^* mice with prion seeding in spinal cord was compared using non-parametric Fisher’s exact tests. For all analyses, p < 0.05 was considered significant.

## Acknowledgements

We thank Biswa Choudhury and Mousumi Paulchakrabarti at the UC San Diego GlycoAnalytics Core for outstanding technical support and mass spectrometry analysis, and the animal care staff at UC San Diego for excellent animal care. We thank Daniel Ojeda-Juárez for critical review of the manuscript. This study was supported by the National Institutes of Health grants NS069566 (CJS), NS076896 (CJS), AG061251 (PAC), the CJD Foundation (CJS), and the Ramón Areces Foundation (PAC). This work was supported in part by the Intramural Research Program of the NIAID (BC and CDO), the Swedish Research Council grant 2016-00748 (KPRN); Project 3 of NHLBI grant HL131474 (JDE) and NSF grant IOS- 2031989 (JDE).

## Notes

The authors have declared that no conflict of interest exists.

### Competing Interest Statement

The authors have declared no competing interest.

